# Modulation of global transcriptional regulatory networks as a strategy for increasing kanamycin resistance of EF-G mutants

**DOI:** 10.1101/059501

**Authors:** Aalap Mogre, Aswin Sai Narain Seshasayee

## Abstract

Evolve and resequence experiments have provided us a tool to understand bacterial adaptation to antibiotics by the gain of genomic mutations. In our previous work, we used short term evolution to isolate mutants resistant to the ribosome targeting antibiotic kanamycin. We had reported gain of resistance to kanamycin via multiple different point mutations in the translation elongation factor G (EF-G). Furthermore, we had shown that the resistance of EF-G mutants could be increased by second site mutations in the genes *rpoD* / *cpxA* / *topA* / *cyaA.* Mutations in three of these genes had been discovered in earlier screens for aminoglycoside resistance. In this work we expand on our understanding of these second site mutations. Using genetic tools we asked how mutations in the cell envelope stress sensor kinase (CpxA^F218Y^) and adenylate cyclase (CyaA^N600Y^) could alter their activities to result in resistance. We found that the mutation in *cpxA* most likely results in an active Cpx stress response. Further evolution of an EF-G mutant in a higher concentration of kanamycin than what was used in our previous experiments identified the *cpxA* locus as a primary target for a significant increase in resistance. The mutation in *cyaA* results in a loss of catalytic activity and probably results in resistance via altered CRP function. Despite a reduction in cAMP levels, the CyaA^N600Y^ mutant has a transcriptome indicative of increased CRP activity, pointing to an unknown non-catalytic role for CyaA in gene expression. From the transcriptomes of double and single mutants we describe the epistasis between the mutation in EF-G and these second site mutations. We show that the large scale transcriptomic changes in the topoisomerase I (FusA^A608E^-TopA^S180L^) mutant likely result from increased negative supercoiling in the cell. Finally, genes with known roles in aminoglycoside resistance were present among the mis-regulated genes in the mutants.

## Introduction

The efficacy of antibiotics, once heralded as miracle drugs, is now under threat because of the emergence of resistance (Davies, 1996; Levy & Marshall, 2004). One way in which bacteria become resistant to antibiotics is by gaining genomic mutations. These genomic mutations tend to accumulate mostly in the target of the antibiotic, and more often than not result in a fitness defect because these target genes tend to be essential or important for cell growth (Lenski, 1998; Andersson & Levin, 1999; Björkman & Andersson, 2000; Andersson, 2006; Andersson & Hughes, 2010). Resistance can also evolve via mutations in non-target genes (Kern et al., 2000), and studying such mutations will yield insight into the mechanism of action of the antibiotic inside the bacterial cell.

Aminoglycosides, a group of ribosome targeting antibiotics (Becker & Cooper, 2013), have a target that is difficult to modify mutationally. This difficulty arises, especially in fast growing organisms like *E. coli*, because of the presence of multiple copies of the gene encoding the target of aminoglycosides, i.e. the 16S rRNA. On short timescales, it is not possible to mutate all copies of the target gene, seven of which are present in *E. coli* for instance, to achieve resistance (Kotra, Haddad & Mobashery, 2000).

In our previous work, we evolved *E. coli* in different sublethal levels of a model aminoglycoside kanamycin (Mogre et al., 2014). We obtained multiple kanamycin resistant mutants of the translation elongation factor EF-G (FusA). At the lower 4-kan (4 μg/ml; 25% lethal concentration kanamycin), we found a single point mutation in EF-G (FusA^P610T^), whereas at the higher 8-kan (50% lethal concentration kanamycin), we found two different point mutations in EF-G (FusA^A608E^ and FusA^P610L^). The FusA^P610T^ allele dominated evolved populations for five transfers (rounds of subculture) in 4-kan; whereas in 8-kan, the FusA^A608E^ allele appeared in the first round of growth, followed by the FusA^P610L^ allele in the next transfer. Among the three EF-G mutants, the FusA^P610L^ allele had the best growth in 8-kan. Interestingly the FusA^A608E^ allele had also accumulated second site mutations in four genes, viz., *rpoD, cpxA, topA* and *cyaA*, in four different isolates. Apart from our work, evolution experiments in aminoglycosides done by Lazar et al., have revealed resistance conferring mutations in *fusA, rpoD, cpxA* and *crp* (whose protein product acts downstream of CyaA), but not in *cyaA* and *topA* (Lázár et al., 2013).

EF-G is a translation factor and a part of this protein, specifically the tip of domain IV, interacts with the decoding centre, the binding site of aminoglycosides (Feldman et al., 2010). Thus, whereas the contribution of EF-G – a factor associated with the binding site of the antibiotic – to resistance is easier to understand, the mechansims by which these second site mutations confer resistance are not immediately apparent. Interestingly, all of the above second site mutations could affect transcription. More specifically, RpoD is the major sigma factor responsible for much of transcription in exponentially-growing *E. coli* (Feklístov et al., 2014). CpxA is an envelope stress sensor kinase, which by phosphorylating its response regulator CpxR, activates the expression of genes that tackle membrane stress (Hunke, Keller & Müller, 2012). Activation of the Cpx response upon antibiotic exposure was thought to result in increased oxidative stress and consequently cell death (Kohanski et al., 2008). However, more recent work has shown that Cpx activation confers resistance and not sensitivity to certain antibiotics (Mahoney & Silhavy, 2013; Manoil, 2013). Topoisomerase I (TopA) relaxes negatively supercoiled DNA, and can thereby affect the transcription of many genes (Travers & Muskhelishvili, 2005). Adenylate cyclase (CyaA) produces cAMP (cyclic adenosine monophosphate), a cellular second messenger, that can influence the expression of a large number of genes via the global regulator CRP (cAMP Receptor Protein) (Zheng et al., 2004). Furthermore, Girgis et al., were able to show that disruptions of *cyaA* and *crp* were beneficial to growth in aminoglycosides (Girgis, Hottes & Tavazoie, 2009).

To understand the contribution of these second site mutations to kanamycin resistance, we first generated their single mutant versions. We found that the second site mutations by themselves provide only a marginal increase in growth in kanamycin. These second site mutations however allow better growth in kanamycin in either the FusA^P610T^ or the FusA^A608E^ background. By comparing these second site mutants with their corresponding whole gene deletions, we attempted to clarify their roles in kanamycin resistance. Further, using RNA-seq, we found a non-additive effect between the mutation in EF-G and that in the second-site on gene expression. By comparing our transcriptome data with previously published datasets, alongwith measurements of plasmid supercoiling, we provide tentative evidence for elevated negative supercoiling in the chromosome of the FusA^A608E^-TopA^S180L^ mutant. Lastly, we were able to see sets of genes with known roles in aminoglycoside resistance among those mis-regulated in these mutants. This reinforces the idea that mutating promiscuous regulators of transcription might be an effective early strategy for adaptation to stress (Finkel, 2006; Wang et al., 2010).

## Materials & Methods

### Strain construction

All strains used had the non-pathogenic *E. coli* K12 MG1655 background. RpoD^L261Q^, CpxA^F218Y^ and CyaA^N600Y^ mutants were constructed from their respective FusA^A608E^-RpoD^L261Q^ / CpxA^F218Y^ / CyaA^N600Y^ double mutants by replacing the FusA^A608E^ allele with the wildtype *fusA* allele linked to a kanamycin resistance cassette. This was done by P1 phage transduction according to the Court lab protocol (Thomason, Costantino & Court, 2007). Selection with a higher concentration of kanamycin (70-80 μg/ml) ensured that only transductants with the wildtype *fusA* linked to the kanamycin resistance cassette were selected, and that the non-transduced recipient double mutants were not. Background growth of the non-transduced double mutants was a common problem; however only the best growing colonies were picked since these would contain the kanamycin resistance cassette. All transductants were verified by PCR to ensure the presence of the kanamycin cassette. The kanamycin resistance cassette was flanked by FRT sites and thus was flipped out using the site specific recombinase Flp provided by the plasmid pCP20. Finally the temperature sensitive plasmid pCP20 was cured from these cells by growing them at 42 ^0^C. The wildtype strain containing the kanamycin cassette near *fusA* was also treated similarly to generate a strain containing the FRT site near *fusA* (WTfrt) and was used as the reference strain. The mutations in *fusA, rpoD, cpxA, topA* and *cyaA* were checked by Sanger sequencing.

Knockout strains Δ*cyaA::kan^R^, Acrp::kan^R^* were earlier generated in the lab, whereas Δ*cpxA::kan^R^* and *AcpxR::kan^R^* were obtained from Coli Genetic Stock Center (CGSC). These knockouts were transferred into the wildtype used in this study, FusA^P610T^ and FusA^A608E^ strains by phage transduction. The process outlined above was used to get rid of the kanamycin resistance cassette after transferring the gene knockouts to the relevant strains including the wildtype used in this study.

### Growth curves and Minimum Inhibitory Concentration (MIC) determination

Growth curves were performed in Luria-Bertani broth (LB) in either flasks or 96-well plates. For the purpose of sample collection and RNA extraction growth curves were performed in flasks at 37 Oc, 200 rpm with optical density readings measured at 600 nm using a Metertech SP-8001 Spectrophotometer. For strain comparisons, growth curves were performed in 96-well plates. These growth curves were performed using the Tecan Infinite F200pro plate reader. The machine incubated the plate at 37 °C and carried out shaking at 198 rpm with optical density readings measured at 600 nm every 15 minutes.

MICs were measured as previously described, using a modification of the broth dilution technique (Mogre et al., 2014).

### Cyclic adenosine monophosphate (cAMP) estimation

Estimation of intracellular cAMP levels was carried out using the cyclic AMP Select EIA kit (501040, Cayman Chemical). Cells growing exponentially (~1.5 hours in LB) and in the stationary phase (~12-15 hours in LB) were harvested by centrifugation at 13,000 g for 1 minute. Cells were immediately transferred onto ice to prevent breakdown of cAMP by phosphodiesterases. Cell pellets were washed once with TBST (20 mM Tris, 150 mM NaCl, 0.05% Tween 20, pH 7.5) before being resuspended in 0.05 N HCl. Cells were then boiled for 5 minutes to extract cAMP. Cells were then spun down at 14000 g and the supernatant containing cAMP was collected. Estimations of cAMP were carried out according to the kit's instructions with the exception that the provided cAMP standard was diluted in 0.05 N HCl to generate the standard curve, since HCl was used for the extraction process.

### Chloroquine gel analysis

Overnight grown cultures of wildtype, Fus^P610T^, Fus^A608E^ and Fus^A608E^-TopA^S180L^, transformed with the pUC18 plasmid, were diluted 1:100 in LB and grown to exponential and stationary phase. pUC18 extraction was done using the QIAprep Spin Miniprep Kit (Qiagen). About 1 μg plasmid was loaded onto 1% agarose gel containing 2.5 μg/ml chloroquine made in 2X Tris Borate EDTA (TBE) buffer. Samples were run in 2X TBE at 3V/cm for 17 hours. After the run, the gel was washed with tap water for three hours, with the water being changed every 20 minutes to remove chloroquine. After the wash, the gel was stained with 1 μg/ml ethidium bromide solution for 1 hour. After a brief wash with tap water for a minute to get rid of excess stain, the gel was illuminated with UV to visualize the different plasmid topoisomers. At the concentration of chloroquine used in these experiments, more supercoiled forms of the plasmid migrate further in the gel. Two biological replicates were analyzed for each strain and the results of one are shown.

### Evolution in kanamycin

The MIC of kanamycin of the FusA^P610T^ mutant was around 60 μg/ml. 25% of this concentration, i.e. 15 μg/ml, was selected for evolving FusA^P610T^ towards higher resistance. Evolution experiments were carried out by batch transfers in LB with and without kanamycin. Two overnight grown replicate populations of *E. coli* were diluted 1:100 in 100 ml LB containing 15 μg/ml kanamycin and 100 ml plain LB as control. Thus two populations evolving in 15 μg/ml kanamycin and two control populations evolving in plain LB were incubated at 37 Oc, 200 rpm for 24 hours before the next transfer. Each evolving population was transferred by 1:100 dilution into fresh medium. The concentration of kanamycin was not changed during the course of the evolution experiment. MICs of all populations were followed at the end of each transfer.

Genomic DNA was extracted from both control and experimental populations using the GenElute Bacterial Genomic DNA Kit (NA2120, Sigma-Aldrich). Integrity of the extracted gDNA was checked on agarose gel, and quality and concentration were checked using NanoDrop UV-Vis Spectrophotometer (Thermo Scientific). Library preparation for deep sequencing was carried out using the Truseq Nano DNA Library Preparation Kit (FC-121-4001, Illumina). Paired end sequencing was carried out using the Illumina HiSeq sequencer (2 x 100 Cycles) at the Next Generation Genomics Facility, Centre for Cellular and Molecular Platforms (C-CAMP). FASTX (http://hannonlab.cshl.edu/fastx_toolkit/) quality filtered reads were trimmed using CUTADAPT version 1.9.dev1 (Martin, 2011), to remove adapter sequences. Error tolerance in identifying adapters was set to 20% and trimmed reads with less than 30 bases were discarded. These trimmed reads, were then mapped to the *E. coli* K12 MG1655 reference genome (NC_000913.3) using BWA mem version 0.7.5a-r405 (Li & Durbin, 2010); paired files were input together at this step. SAMTOOLS version 1.3 (Li et al., 2009) was then used to generate the pileup file from the sam files generated by BWA. Finally, the list of single nucleotide polymorphisms (SNPs) and indels was compiled from the pileup file using VARSCAN version 2.3.8 (Koboldt et al., 2012).

### RNA extraction, sequencing and analysis

For RNA extraction, cells were grown in LB and harvested at the point of maximal growth rate (Fig. S1) after the addition of stop solution to stabilise cellular RNA and stop transcription. Two biological replicates were harvested for each strain including the reference strain WTfrt. RNA was extracted using the hot phenol-chloroform method. DNase treated RNA was depleted of ribosomal RNA using the Ambion Microbe Express Kit (AM1905). RNA was checked for quality using Bioanalyzer (Agilent). Checked RNA was used for library preparation and sequencing. RNA quality checks, library preparation and sequencing were carried out at Genotypic (India). Briefly, 100 ng of qubit quantified RNA was used for library preparation using the NEXTflex Rapid Directional RNA-Seq kit (5138-08 Bioo Scientific). The library was quantified using qubit and its quality was checked using Agilent Bioanalyzer before proceeding for sequencing on the Illumina NextSeq 500 sequencer.

FASTX filtered reads were trimmed using CUTADAPT and aligned to the *E. coli* reference genome (NC_000913.3) using BWA. SNP and indel calling was done to ensure that the correct mutations were present in the relevant samples (Fig. S2). The number of reads mapping to each gene was obtained using custom Python scripts. Correlation of raw read counts between replicates were high (>0.9, Fig. S3). Even across different strains, the strength of correlation was high (>0.8, Fig. S3). We also checked that genes within operons were similarly expressed (Fig. S4).

Subsequently the R (R Core Team, 2016) package EdgeR (Robinson, McCarthy & Smyth, 2010) was used to call differentially expressed genes using a P value cutoff of 0.001 (using the Benjamini Hochberg method to control the false discovery rate in multiple testing).

Gene ontology analysis was carried out using the R package topGO (Alexa & Rahnenfuhrer, 2010). *E. coli* gene annotations were obtained from Ecocyc (Karp et al., 2014) and gene ontology terms were obtained from the Gene Ontology Consortium (Gene Ontology Consortium, 2015). In topGO, the Fisher test was used to assess significance of enriched gene sets and terms with P values < 0.01 were considered significant.

All scripts are available on Github (https://github.com/aswinsainarain/mogre_kan_secondsite), as well as on http://bugbears.ncbs.res.in/mogre_kan_secondsite/.

## Results and Discussion

### Second site mutations increase kanamycin resistance of EF-G mutants

In our previous work we had found that four 'second-site’ mutations, namely RpoD^L26lQ^, CpxA^F2l8Y^, CyaA^N600Y^ and TopA^S180L^, appeared on a FusA^A608E^ background in 8-kan. The double mutants had marginally greater resistance to kanamycin, as measured by their MICs, than the single FusA^P610T/A608E^ mutants; this difference was statistically significant in the case of the FusA^A608E^ mutant, but not the FusA^P610T^ mutant (Fig. 1A). However, the double mutants grew to a higher cell density in 8-kan, thus pointing to their selective advantage over the single mutant in kanamycin (Fig. 1B).

**Figure 1:**
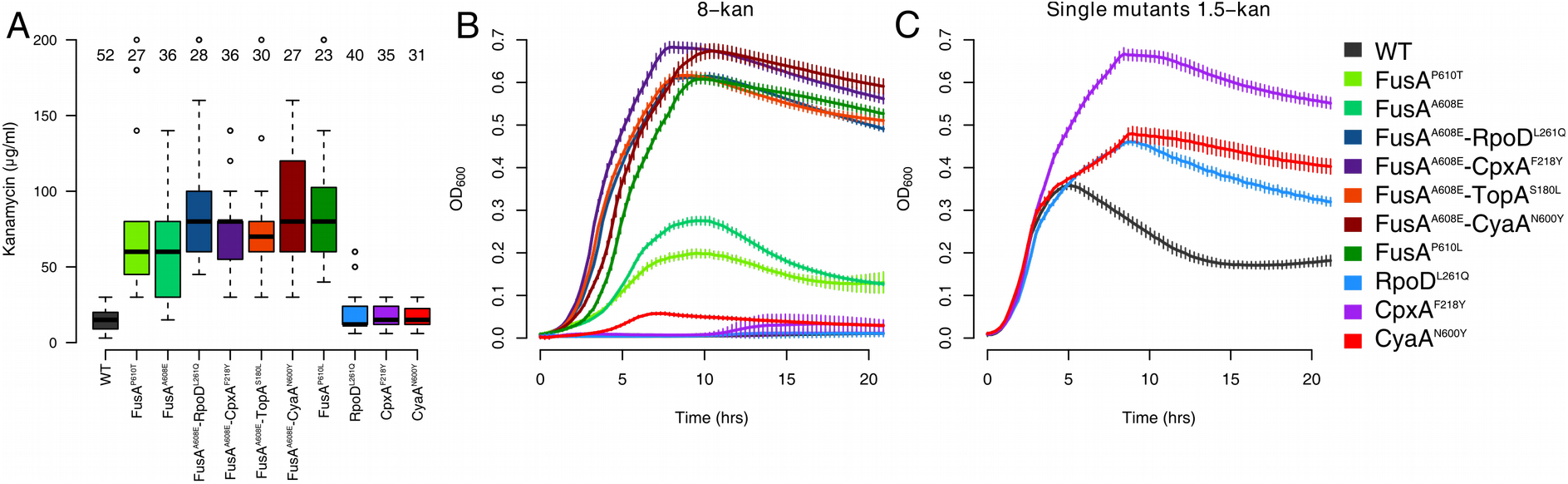
**Kanamycin resistance of mutants.** (A) Boxplots showing distributions of Minimal Inhibitory Concentrations (MICs) of kanamycin of the wildtype (WT) and various mutants. The number of replicates is mentioned over each boxplot. All mutants, except RpoD^L261Q^, CpxA^F218Y^ and CyaA^N600Y^, are significantly more resistant than the wildtype (Welch two sample t-test, P value < 10^-7^). Although the medians of the FusA^A608E^-RpoD^L261Q^ / CpxA^F218Y^ / TopA^S180L^ / CyaA^N600Y^ double mutants tend to be higher than that of the FusA^P610T^ mutant, this difference is not statistically significant (P value > 0.09). The difference between the medians of these double mutants and the FusA^A608E^ mutant are statistically significant (P value < 0.02), except in the case of the FusA^A608E^-TopA^S180L^ mutant (P value 0.061) (B) Growth of mutants in 8-kan (8 μg/ml kanamycin). (C) Growth of the RpoD^L261Q^, CpxA^F218Y^ and CyaA^N600Y^ mutants in 1.5-kan (1.5 μg/ml kanamycin). In B and C, error bars represent standard deviation of eight replicates.

We replaced the FusA^A608E^ mutation in all the double mutants with the FusA^P610T^ mutation that had evolved in the wildtype background in 4-kan. The FusA^P610T^ mutation showed sub-optimal growth under conditions in which FusA^A608E^ and its second-site mutants had emerged (8-kan, Fig. 1B). These FusA^P610T^ double mutants grew to a higher cell density than the FusA^P610T^ single mutant alone in 8-kan (Fig. S5, control growth curves in 0-kan in Fig. S6). Thus, the selective advantage conferred by these second site mutations was similar between the two primary FusA mutants.

We constructed single mutant versions of RpoD^L261Q^, CpxA^F218Y^ and CyaA^N600Y^ from their respective double mutants by replacing the mutant FusA^A608E^ allele with the wildtype allele (see Materials & Methods). For reasons not understood, we were unable to construct the TopA^S180L^ single mutant. We saw that the resistance of the double mutants decreased to almost wildtype levels in these single second-site mutants (Fig. 1A). This is seen more clearly in the growth curves of these single mutants in 8-kan (Fig. 1B). These single mutants, however, fared better than the wildtype at a very low concentration of kanamycin (Fig. 1C).

From the order of occurrence of mutations in our evolution experiments, and the inability of single mutants of *cpxA, cyaA* and *rpoD* to grow well in kanamycin, it is reasonable to suggest that the mutation in EF-G potentiates the second site mutations, which further increase its resistance.

### Mutation in the extra-cytoplasmic stress sensor CpxA results in resistance via hyper-activation of the Cpx stress response

The Cpx stress response is mediated by the CpxA sensor kinase and its cognate response regulator CpxR (~58 targets in RegulonDB). This versatile two-component system responds to various kinds of stress signals, especially those associated with membrane stress (Pogliano et al., 1997; Raivio & Silhavy, 1997; Hunke, Keller & Müller, 2012; Vogt & Raivio, 2012). Aminoglycosides cause the accumulation of misfolded proteins in the cell membrane and periplasmic space (Bryan & Kwan, 1983; Davis, 1987; Kohanski et al., 2008). The Cpx system responds to this stress (Kohanski et al., 2008). It was thought that the crosstalk of the Cpx response with the redox reactive Arc two-component system results in oxidative stress that kills cells (Kohanski et al., 2008). However, more recent genetic experiments have revealed that activation of the Cpx response has a protective role (Mahoney & Silhavy, 2013).

To understand the effect of the point mutation in *cpxA* on the activity of the Cpx response, we deleted *cpxR* in the EF-G mutants and the wildtype to see whether this increased or decreased their growth in kanamycin. Deleting the response regulator CpxR and not the sensor kinase CpxA is the best way to attenuate the Cpx response. This is because, in the absence of CpxA, CpxR can be cross-activated by other kinases, and this can in fact result in the hyper-activation of the Cpx response. FusA^A608E^-Δ*cpx*R resulted in a decrease in growth in kanamycin, while there was no discernible effect of the deletion on the wildtype (Fig. 2A). This suggests that an intact Cpx response is essential for the resistance of these mutants.

**Figure 2:**
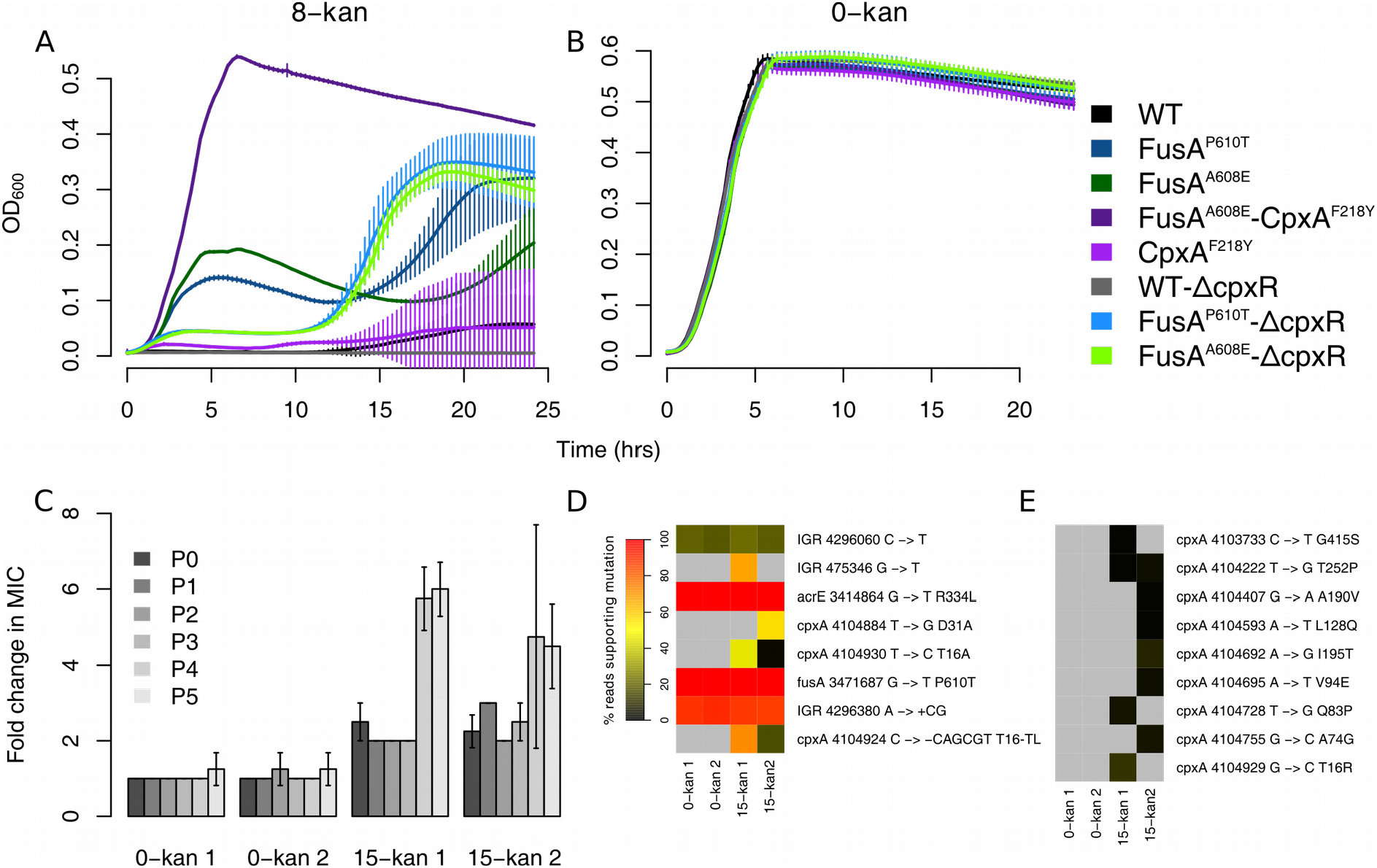
**Activation of the Cpx response results in resistance.** Growth curves in 8-kan (A) and 0-kan (B). The labels for the x and y axis are common. Plotted are the means from eight replicates with error-bars representing standard deviation. In the 8-kan growth curves the huge error bars in some of the strains are produced when a few replicates start growing, possibly due to acquisition of some resistance conferring mutation, and thus this error cannot be eliminated. (C) Fold changes in MICs of populations evolved in 15-kan over the MICs of populations evolved in 0-kan. Two replicate populations grown in either 0-kan or 15-kan are shown. MICs of evolving populations at the end of growth (24 hours) after each batch transfer were determined and are represented by P0-P5. The MIC of the 0-kan 1 population at P0 was used to calculate fold changes. Error bars represent standard deviation of four replicates. (D) Heatmaps showing the abundance of variants revealed by sequencing of both control and evolved populations. The colour represents the percentage of reads supporting mutations and is an approximate proxy for the abundance of the mutation. The list of variants was trimmed such that only mutations present in greater than 20% of the reads in at least one sample were retained. (E) Heatmaps showing abundance of all low-frequency cpxA variants. The list of variants was trimmed to include only mutations in cpxA. The mutations shown in (D) are not shown here. The colour scale is as shown in (D).

We saw that the FusA^A608E^-Δ*cpx*R mutants showed a spurt in growth at a certain point in batch culture, whence these mutants grew better than their EF-G single mutant counterparts (Fig. 2A). This was consistent across the different transductants tested (Fig. S7 and S8). However, even after this growth spurt, their saturation optical density never reached that of the FusA^A608E^-CpxA^F218Y^ double mutant obtained from the evolution experiment (Fig. 2A).

Control growth experiments in the absence of kanamycin told us that the decreased growth of the FusA^A608E^-Δ*cpx*R strains seen in kanamycin was not the result of any generic growth defect conferred by the deletion itself (Fig. 2B). Thus, it is reasonable to conclude that the Cpx system is active, and perhaps hyperactive, in the double mutant, and that its activity is linked to resistance. CpxR has fifty eight targets in RegulonDB (Gama-Castro et al., 2016). However only a handful of these genes are among the differentially regulated genes in the FusA^A608E^-CpxA^F218Y^ and CpxA^F218Y^ mutants in relation to the wildtype (18 in the CpxA^F218Y^ mutant and 10 in the FusA^A608E^-CpxA^F218Y^ mutant), as measured by RNA-seq experiments of these mutants in the absence of kanamycin. There are nearly equal numbers of positive and negative targets of CpxR among the up and down-regulated genes. Thus, the status of the Cpx response is not clarified by a bird's eye view of the transcriptome of these mutants. However, *cpxA* and *cpxP* are up-regulated in both the FusA^A608E^-CpxA^F218Y^ and CpxA^F218Y^ mutants; and *cpxR* is up-regulated in the CpxA^F218Y^ mutant. All these three genes are positive targets of CpxR, and their up-regulation in the mutant suggests that the Cpx system might be hyper-active.

### Further evolution of the FusA^P610T^ mutant in kanamycin reveals *cpxA* as the primary locus targeted for a further increase in resistance

In our previous evolution experiment, while the FusA^A608E^ mutant rapidly accumulated second site mutations in 8-kan, the FusA^P610T^ mutant which had evolved in 4-kan did not accumulate other mutations even after five transfers (Mogre et al., 2014). Thus we decided to evolve FusA^P610T^ in a higher concentration of kanamycin (15-kan; 15 μg/ml kanamycin; 25% of the MIC of FusA^P610T^) to see what second site mutations would accumulate and eventually dominate in the FusA^P610T^ background. The evolution experiment involved serial batch transfers of pure FusA^P610T^ mutant populations in 15-kan every twenty four hours. MICs of all populations were followed at the end of each transfer throughout the experiment.

We observed an initial increase in MIC of around two-fold almost immediately and it notched up further at the end of the fourth transfer to around five to six-fold (Fig. 2C). At the end of the fifth transfer, genomic DNA from both control and evolved populations were sequenced. We looked at the mutations that were present in at least one sample in greater than 20% frequency (Fig. 2D). In the populations evolved in 15-kan we saw multiple mutations in *cpxA* present in different frequencies. There were two point mutations and one deletion of 6 nucleotides in *cpxA.* We also found several other cpxA mutations, present at very low frequencies (<20%), only in the populations exposed to kanamycin (Fig. 2E). Thus we see a heterogeneous population with different mutations in *cpxA.* This heterogeneity might be responsible for the large variations in the MIC determinations of these populations (Fig. 2C).

Residue changes in CpxA that confer kanamycin resistance were scattered across the protein and were located in helix-I, periplasmic and cytoplasmic-II domains (Fig. S9). In particular, mutations in helix-I reached high frequencies (>50%) in the FusA^P610T^ populations evolved in 15-kan. The residue T16 in helix-I was particularly targeted with three different mutations, two of which reached high frequencies. Mutations in the periplasmic domain, helix-II and cytoplasmic-II domain are known to result in kanamycin resistance due to hyperactivation of CpxA (Raivio & Silhavy, 1997). However, we did not see any mutations in helix-II.

We noticed that multiple low frequency mutations in the gene *sbmA* had appeared in the populations evolved in kanamycin (Fig. S10). The product of this gene is involved in the transport of peptide antibiotics and its deletion results in increase of resistance to antimicrobial peptides (Laviña, Pugsley & Moreno, 1986; Yorgey et al., 1994; Salomón & Farías, 1995; Saier et al., 2009; Corbalan et al., 2013; Runti et al., 2013; Paulsen et al., 2016).

To summarize, the *cpxA* locus seems to be the primary region targeted for the next significant increase in resistance of the EF-G point mutant.

### Disruption of adenylate cyclase catalytic activity gives kanamycin resistance mediated by altered CRP function

Adenylate cyclase is an enzyme that catalyzes the synthesis of cyclic adenosine monophosphate (cAMP) from ATP. cAMP functions as a second messenger in *E. coli* (Botsford & Harman, 1992; McDonough & Rodriguez, 2012). A well-known mechanism by which cAMP alters gene expression is by binding to and allosterically activating the global transcription regulator cAMP receptor protein (CRP) (Botsford & Harman, 1992; McDonough & Rodriguez, 2012), which has 477 targets in RegulonDB. Girgis et al., subsequent to a transposon mutagenesis screen, demonstrated that deletions of *cyaA* and *crp* increased resistance to aminoglycosides (Girgis, Hottes & Tavazoie, 2009). Furthermore, inactivation of adenylate cyclase was shown to result in activation of the Cpx system (Strozen, Langen & Howard, 2005) which has a known role to play in kanamycin resistance (Mahoney & Silhavy, 2013).

We found that the level of cAMP in both the CyaA^N600Y^ and FusA^A608E^-CyaA^N600Y^ mutants was lower than that in the wildtype and comparable to that in *AcyaA* (Fig. 3A). Thus, the CyaA^N600Y^ mutation results in a loss of catalytic activity. The greater spread in the cAMP levels in the CyaA^N600Y^ and FusA^A608E^-CyaA^N600Y^ mutants suggests that the mutated adenylate cyclase might have reduced catalytic activity, and not a complete loss of function. This is consistent with the fact that this mutation is present in the regulatory domain of the protein and not the catalytic domain.

**Figure 3:**
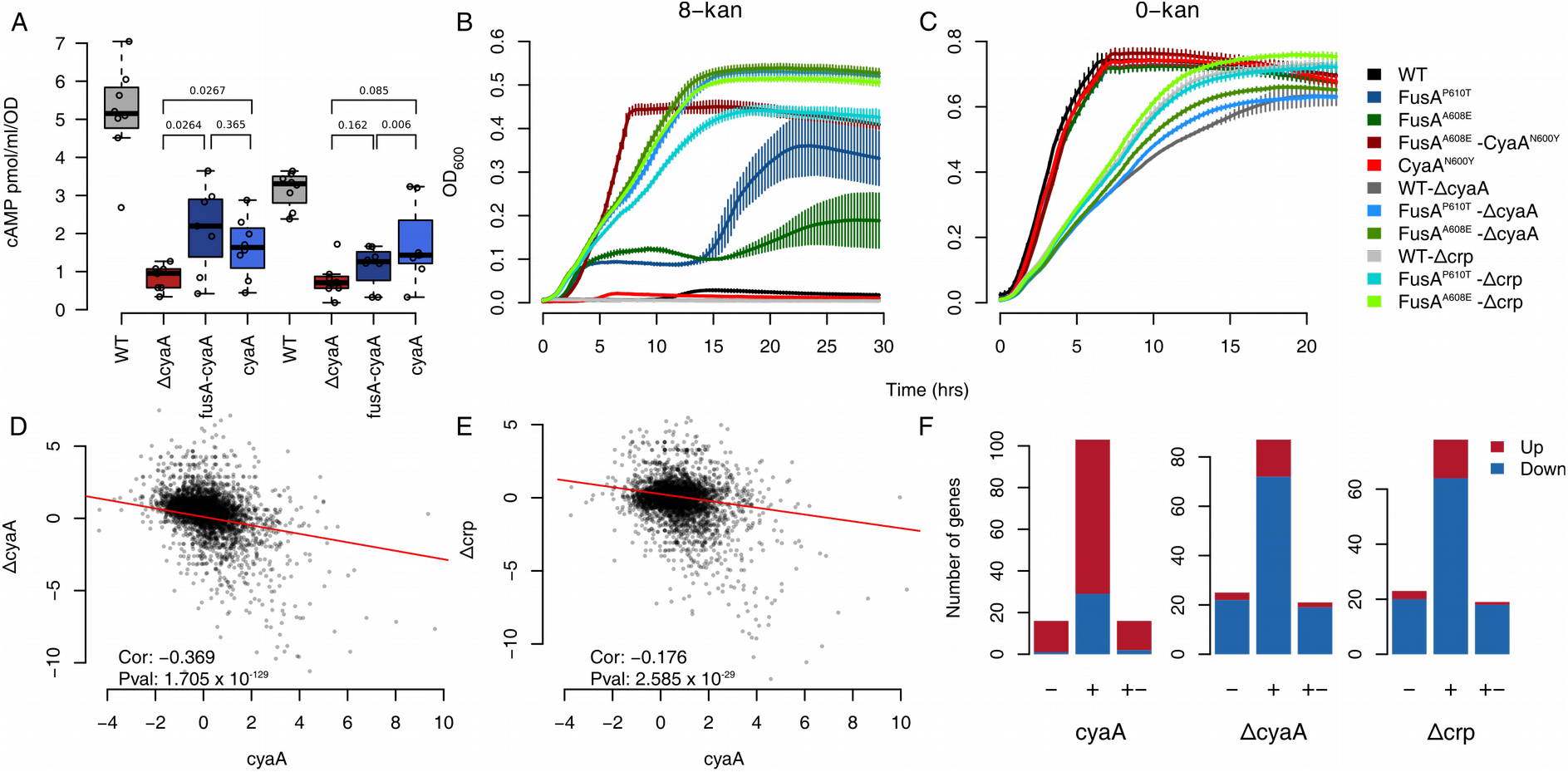
**Inactivation of adenylate cyclase results in resistance.** (A) Boxplots showing the distribution of cAMP estimates of strains in the exponential and stationary phase. The difference between the wildtype and the other mutants are significant (P value < 1 x 10^-2^). P values for other relevant comparisons are mentioned in the plot. The FusA^A608E^-CyaA^N600Y^ mutant is referred to as fusA-cyaA and the CyaA^N600Y^ mutant as cyaA. (B & C) Growth curves in 8-kan (B) and 0-kan (C). The labels for the x and y axis are common. Plotted are the means from eight replicates with error-bars representing standard deviation. In the 8-kan growth curves the huge error bars in some of the strains are produced when a few replicates start growing, possibly due to acquisition of some resistance conferring mutation, and thus this error cannot be eliminated. (D-E) Scatter plots comparing Log2 fold-changes of genes in the CyaA^N600Y^ mutant with those in the ΔcyaA (D) /Δcrp (E) knockout strains. The mutant is referred to by its gene name for brevity. The time-points for cell harvesting for RNA extraction of the ΔcyaA / Δcrp strains were similar to that of the mutants. The Spearman correlation coefficient and its P value are mentioned. (C) Barplots showing the the number of targets of CRP present among the up and down-regulated genes in the CyaA^N600Y^, ΔcyaA and Δcrp strains. The numbers of positive (+), negative (-) and dual targets (+-) of CRP present among the up (red) and down-regulated (blue) genes are shown in the stacked barplots. Similar results are seen with the FusA^A608E^-CyaA^N600Y^ mutant and are shown in Fig. S13.

We also found an increase in the stationary phase cell density of the FusA^P610T/A608E^-Δ*cyaA* strains in kanamycin whereas *WT*-Δ*cyaA* was not affected considerably (Fig. 3B). FusA^P610T/A608E^-Δ*crp* also grew to a higher stationary phase cell density in kanamycin, whereas the WT-Δ*crp* did not (Fig. 3B). Both these observations were consistent across multiple transductants tested (Fig. S11).

Both *ΔcyaA* and *Δcrp*, in the FusA^P610T/A608E^ mutant backgrounds, had lower growth rates in kanamycin compared to FusA^P610T/A608E^ and FusA^A608E^-CyaA^N600Y^. The presence of a similar growth defect in the wildtype strain containing these gene knockouts, during growth in the absence of kanamycin, indicated that the growth defect was specific to the gene knockouts (Fig. 3C and S12), and not the mutants isolated.

Together, we conclude that the evolved point mutation in *cyaA* results in kanamycin resistance via an altered adenylate cyclase and subsequently altered CRP function. This mutation has the added benefit of not conferring the growth defect associated with the knockouts of either of these genes.

We next compared the transcriptomes of the CyaA^N600Y^ mutant with the transcriptomes of the *ΔcyaA* and *Δcrp* strains. Surprisingly, fold changes of genes in CyaA^N600Y^, in comparison to the wildtype, negatively correlated with those in the *ΔcyaA* and *Δcrp* strains (Fig. 3D and E). This correlation was low but significant. This is consistent with the results of our comparison with the list of CRP targets available in the RegulonDB database. As expected, genes that are activated by cAMP-CRP are down-regulated in the *ΔcyaA* and *Δcrp* strains (Fig. 3F). However such genes are mostly up-regulated in the CyaA^N600Y^ mutant (Fig. 6C). This stands for the FusA^A608E^-CyaA^N600Y^ mutant as well (Fig. S13).

Thus although the levels of cAMP are low in the CyaA^N600Y^ mutant (Fig. 3A), their transcriptomes are opposite to that of *ΔcyaA* / *Δcrp* (Fig. 3D and E). This is contradictory to expectation, more so since since the FusA^P610T/A608E^-Δ*cyaA* / *Δcrp* strains grow better in kanamycin than the FusA^P610T/A608E^ mutants. It is possible that this phenotypic similarity of the FusA^P610T/A608E^-Δ*cyaA* / *Δcrp* strains to the FusA^P610T^-CyaA^N600Y^ mutant is coincidental and that the full knockout and the point mutant confer resistance through distinct means. We ensured the absence of other mutations in the genomes of the *ΔcyaA* / *Δcrp* transductants using whole genome sequencing (Fig. S14) and the absence of other mutations in the FusA^A608E^-CyaA^N600Y^ and CyaA^N600Y^ strains by calling mutations from the RNA-seq data.

This unexpected behaviour of the CyaA^N600Y^ mutation could stem from the fact that the CyaA^N600Y^ mutation is a point mutation and not a knockout. As a result, the adenylate cyclase protein would still be produced, and this could have alternative functions. There could also be feedback involved: although not called up-regulated, the fold changes of both *cyaA* and *crp* are higher by around 2-fold in the CyaA^N600Y^ and FusA^A608E^-CyaA^N600Y^ mutants (Fig. S15). In line with this, the promoter activities of these two genes were also higher in the CyaA^N600Y^ mutant (Fig. S15A and B), as revealed by promoter-GFP fusions. Thus, further work is required to understand the function of CyaA^N600Y^.

### The FusA^A608E^-TopA^S180L^ mutant displays increased negative supercoiling

The bacterial chromosome is a highly condensed and supercoiled DNA molecule (Woldringh, 2002; Thanbichler, Wang & Shapiro, 2005; Thanbichler & Shapiro, 2006; Toro & Shapiro, 2010). The extent of supercoiling can influence gene expression (Travers & Muskhelishvili, 2005), and is known to be affected by various environmental factors (Rui & Tse-Dinh, 2003) such as osmotic stress (McClellan et al., 1990; Cheung et al., 2003), starvation (Balke & Gralla, 1987), temperature (McClellan et al., 1990) and oxygen tension (Hsieh, Burger & Drlica, 1991). Global supercoiling of the chromosome is maintained by a balance between the activities of enzymes called topoisomerases. DNA gyrase (Topoisomerase II, encoded by the genes *gyrA* and *gyrB)* catalyzes an increase in negative superhelicity. Topoisomerases I *(topA)*, III *(topB)* and IV *(parC* and *parE)* relax negative supercoils (Travers & Muskhelishvili, 2005; Gubaev & Klostermeier, 2014). Topoisomerase IV can also relax positive supercoils, which is its preferred substrate (Crisona et al., 2000) and is involved in decatenation and unknotting of DNA (López et al., 2012). The decatenation activity is triggered by an increase in negative supercoiling (Zechiedrich, Khodursky & Cozzarelli, 1997). The levels of these enzymes in the cell are maintained by a homeostatic feedback from the supercoiled state of the chromosome (Menzel & Gellert, 1987; Tse-Dinh & Beran, 1988; Pruss & Drlica, 1989; Snoep et al., 2002).

One of the second site mutations in the FusA^A608E^ mutant lies in Topoisomerase I (TopA^S180L^). To understand the supercoiling state of the chromosome in the FusA^A608E^-TopA^S180L^ mutant we employed the cholorquine gel assay (Hsieh, Burger & Drlica, 1991) to look at the supercoiling of a reporter plasmid – pUC18 (Fig. 4A). When plasmid DNA is run through an agarose gel containing 2.5 μg/ml chloroquine by electrophoresis, a separation of different topoisomers is achieved, where more supercoiled topoisomers run further than relaxed topoisomers. In both exponential and stationary phases of growth, we found that the pUC18 molecules were more negatively supercoiled, i.e. they ran further on the chlroquine gel, in the FusA^A608E^-TopA^S180L^ mutant than in the wildtype or the single mutants in FusA.

**Figure 4:**
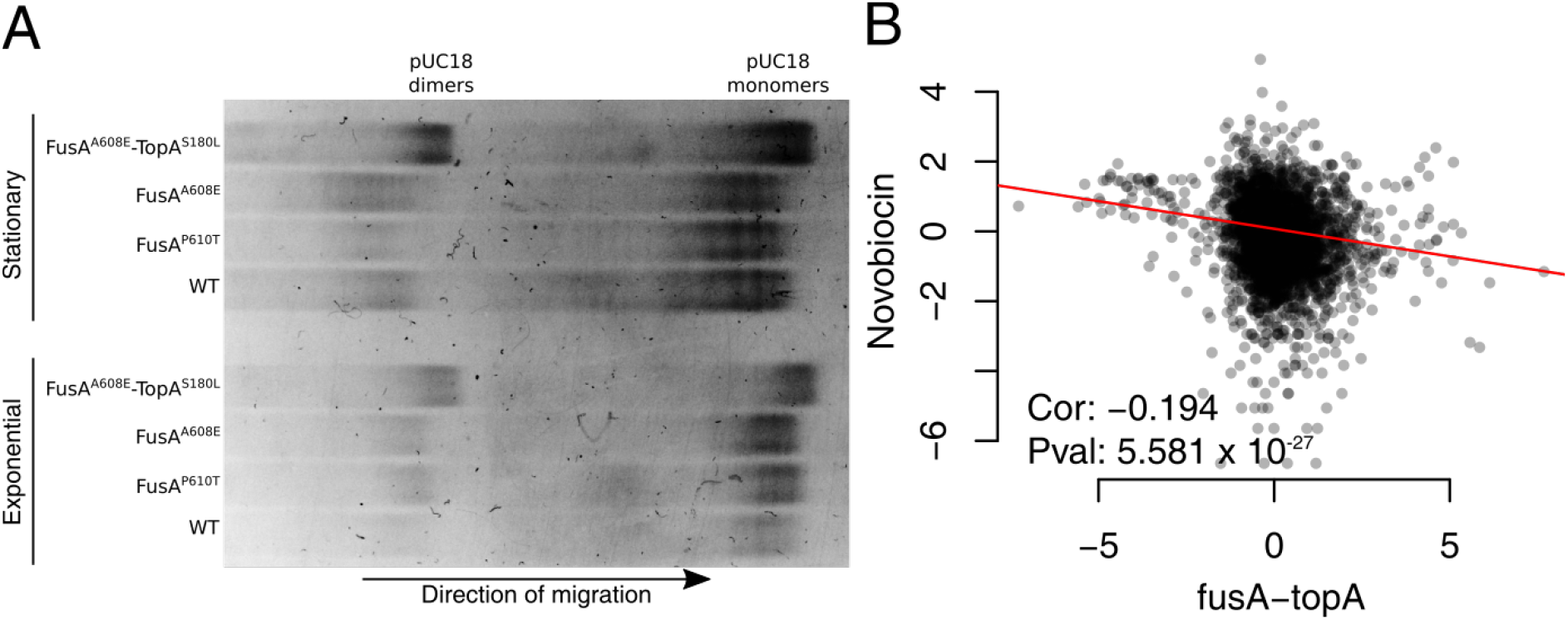
**Evidence for supercoiling changes in the FusA^A60SE^-TopA^S1S0L^ mutant.** (A) Gel picture showing mobility of pUC18 topoisomers on agarose gel containing 2.5 μg/ml chloroquine. (B) Scatter plot showing correlation of fold changes of genes in the FusA^A608E^-TopA^S180L^ mutant with microarray derived gene-expression ratios obtained by inhibiting DNA gyrase function (Peter et al., 2004) using 20 μg/ml novobiocin. For a detailed comparison with the Peter et al. dataset, refer to Fig. S18.

In an earlier gene expression study, Peter et al. used topoisomerase targeting antibiotics and temperature sensitive DNA gyrase mutants to change the supercoiling of the *E. coli* chromosome, and profile the resulting changes in the transcriptome using microarrays (Peter et al., 2004). Their data consists of gene expression ratios obtained from microarrays, loaded with RNA from cells at multiple timepoints after treatment / temperature shift, with RNA from cells before treatment / temperature shift serving as a reference. We found that the gene expression profile of FusA^A608E^-TopA^S180L^ mutant negatively correlates with that of Peter et al's novobiocin treated *E. coli* (Fig. 4B). Negative correlation with novobiocin treatment suggests that there is increased negative supercoiling in the cell, since novobiocin inhibits DNA gyrase. This view is supported by the expression levels of the topoisomerases themselves: as expected from a negative feedback in the presence of high negative supercoiling, the levels of *gyrA* and *gyrB* (DNA gyrase) are low and that of *topA* (Topoisomerase I) is high (GEO accession number GSE82343). Furthermore, we see that up-regulated genes in the mutant are AT rich (Fig S16), which is in accord with the fact that AT rich regions are more susceptible to unwinding in negatively supercoiled DNA. Finally, the FusA^A608E^-TopA^S180L^ mutant shows a slight growth defect at a lower temperature (Fig. S17), which is in line with the known cold sensitivity of the *topA* deletion strain (Stupina & Wang, 2005).

Taken together, we show that the mutation in *topA* reduces its activity and results in increased negative supercoiling in the cell due to intact DNA gyrase function. The loss in activity could also result in increased R-loop formation in the mutant since Topoisomerase I resolves R-loops (Massé & Drolet, 1999; Usongo et al., 2008). Increased translation may help reduce R-loops (Massé & Drolet, 1999; Broccoli et al., 2004; Gowrishankar & Harinarayanan, 2004; Gowrishankar, Leela & Anupama, 2013). This might even potentially explain the possible genetic interaction between *fusA* and *topA.* Increased dosage of genes encoding Topoisomerase IV *(parC* and *parE)* have been shown to relieve growth defects caused by inactivation of *topA* (Kato et al., 1990). We also see an increased expression of *parC* and *parE* in the FusA^A608E^-TopA^S180L^ mutant (GEO Series accession number number GSE82343).

To our knowledge, we are the first to explore the link between chromosomal supercoiling and aminoglycoside resistance. Since we do not have a TopA^S180L^ single mutant we don't understand the effects of the FusA^A608E^ mutation on the TopA^S180L^ mutation. Further experiments will help work out the biochemical activity of the TopA^S180L^ mutant, its genetic interaction with FusA^A608E^, and the mechanism of resistance of this mutant.

### Similarities among mutants

The number of differentially expressed genes varied substantially among the mutants (Fig. 5A). The CpxA^F218Y^, CyaA^N600Y^ and FusA^A608E^-TopA^S180L^ mutants had the most number of differentially expressed genes. The FusA^A608E^, RpoD^L261Q^ and FusA^A608E^-RpoD^L261Q^ mutants had the least number of differentially expressed genes.

**Figure 5:**
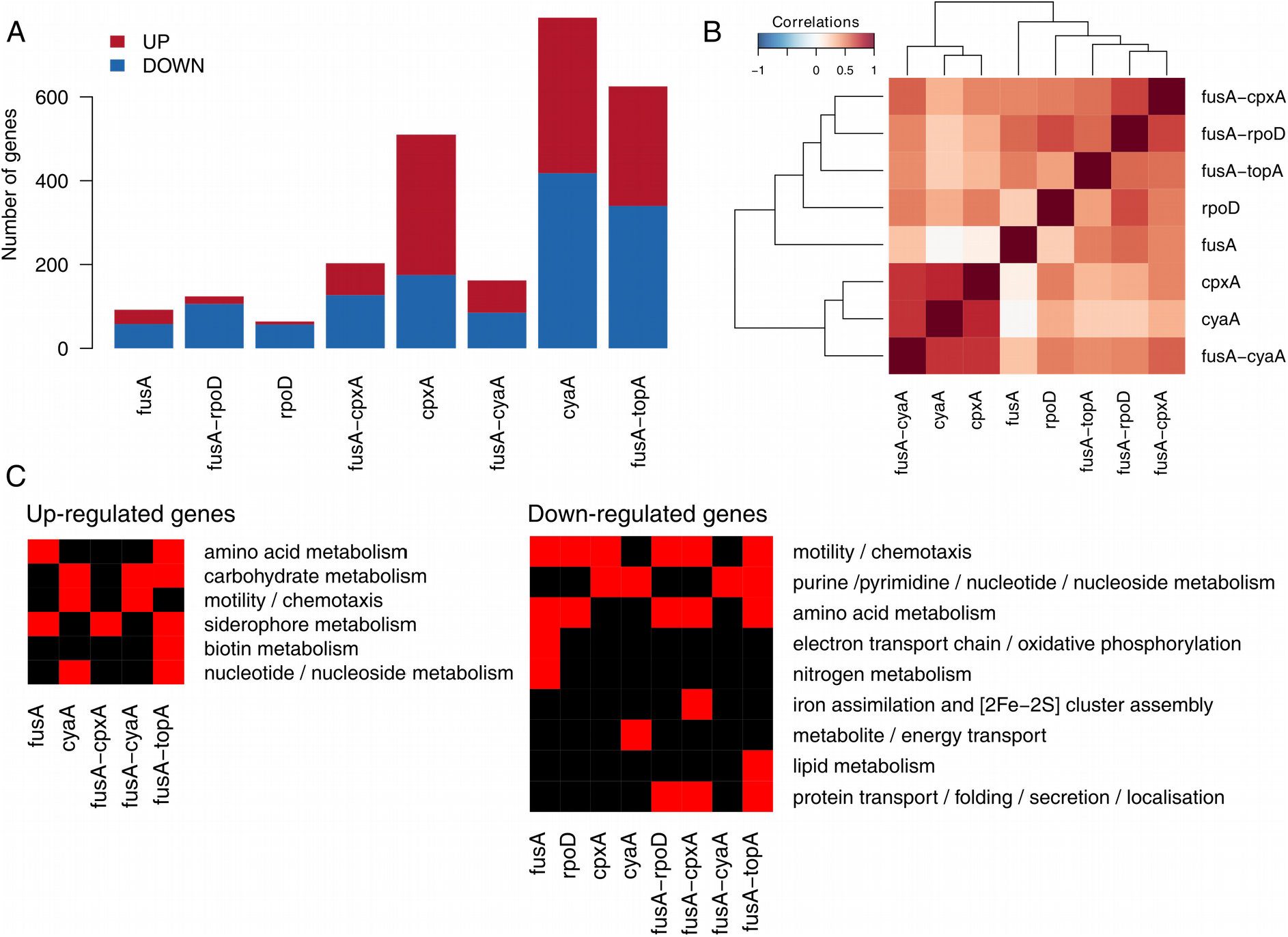
**Summary of differentially expressed genes across mutants.** (A) Numbers of up and down-regulated genes in the mutants. Mutants are referred to by their gene names for brevity. (B) Heatmap showing the matrix of Spearman correlations among mutants. Fold changes of all genes were used to derive these correlations. (C) Heatmap showing enriched gene ontology terms among the up and down-regulated genes in the mutants. Many gene ontology terms have been combined to give this simplified picture. For the entire set of terms and their P values refer to Fig. S20.

We were surprised that the *rpoD* mutants had the least number of differentially expressed genes considering the role of this gene as a transcription initiation factor for around 50% of the genes in *E. coli*, particularly those expressed in exponentially growing cells. How this mutation results in resistance remains elusive. Nonetheless, the location of the mutation is interesting. The mutation resides in a highly conserved residue in the large non-conserved domain of this protein (Fig. S19). While no clear function is ascribed to this domain, certain residues in this domain have been shown to be involved in promoter escape (Leibman & Hochschild, 2007), and one of these residues lies very close to the mutated residue mentioned in our study. It is difficult to hypothesise how a residue involved in allowing the escape of the RNA polymerase from the housekeeping sigma factor to facilitate transcription initiation, must be involved in aminoglycoside resistance (unless this has something to do with coupled transcription and translation), but in the absence of other information remains a valuable lead.

Clustering based on correlation between fold changes of all genes relative to the wildtype across mutants tends to cluster the dataset according to the mutants, but not always so (Fig. 5B). Notably, fold changes of differentially expressed genes in the CyaA^N600Y^ mutant are well correlated with that in the CpxA^F218Y^ mutant. This further highlights the link between CRP and the Cpx response mentioned earlier (Strozen, Langen & Howard, 2005). Thus, as an outcome of the evolution experiment, we see two different mutations resulting in similar transcriptional states. The correlation between the FusA^A608E^-CpxA^F218Y^, FusA^A608E^-RpoD^L261Q^ and FusA^A608E^-TopA^S180L^ mutants were high, offering another example of converging effects of different mutations.

### Effect of second site mutants on gene expression and dependence on EF-G

We evaluated the impact of the FusA^A608E^ mutation on the transcriptomes of the double mutants in detail. The FusA^A608E^ single mutant had roughly thirty genes up and fifty genes down-regulated (Fig. 5A). Sequencing of the transcriptomes of single and double mutants enabled us to look at interactions of the FusA^A608E^ mutant with the second site mutations in terms of fold changes of differentially expressed genes. From the transcriptomes of exponentially growing cells, we saw clear interactions of the primary kanamycin resistance conferring mutation in *fusA* with the second site mutations in *cyaA, cpxA* and *rpoD.*

To assess the extent to which the transcriptomes of the FusA^A608E^ single mutant and CyaA^N600Y^ / CpxA^F218Y^ / RpoD^L261Q^ single mutants explain the gene expression state of the FusA^A608E^-CyaA^N600Y^ / CpxA^F218Y^ / RpoD^L261Q^ double mutants, we plotted the log2 fold change in the double mutant against the sum of the log2 fold changes in the FusA^A608E^ single mutant and the second site single mutants (Fig. S22D, H and L). In the absence of a genetic interaction between the two single mutants, we would expect the scatter plot to lie along the 45° line. We find that this is not the case in each of the three double mutants we evaluated. In other words, the absolute difference in log2 fold change in expression between the double mutant and the sum of the two corresponding single mutants is significantly different from zero (Wilcoxon signed rank test P value < 2.2 x 10^-16^, Fig. S22). These indicate that the FusA^A608E^ background affects the transcriptional state of the CyaA^N600Y^ / CpxA^F218Y^ / RpoD^L261Q^ single mutants in a non-additive manner.

More specifically, we saw that the fold changes of genes differentially expressed in the CyaA^N600Y^ mutant were reduced in the FusA^A608E^-CyaA^N600Y^ mutant (Fig. 6A). Most of these genes were not differentially expressed in the FusA^A608E^ mutant (Fig. 6B and C).

**Figure 6:**
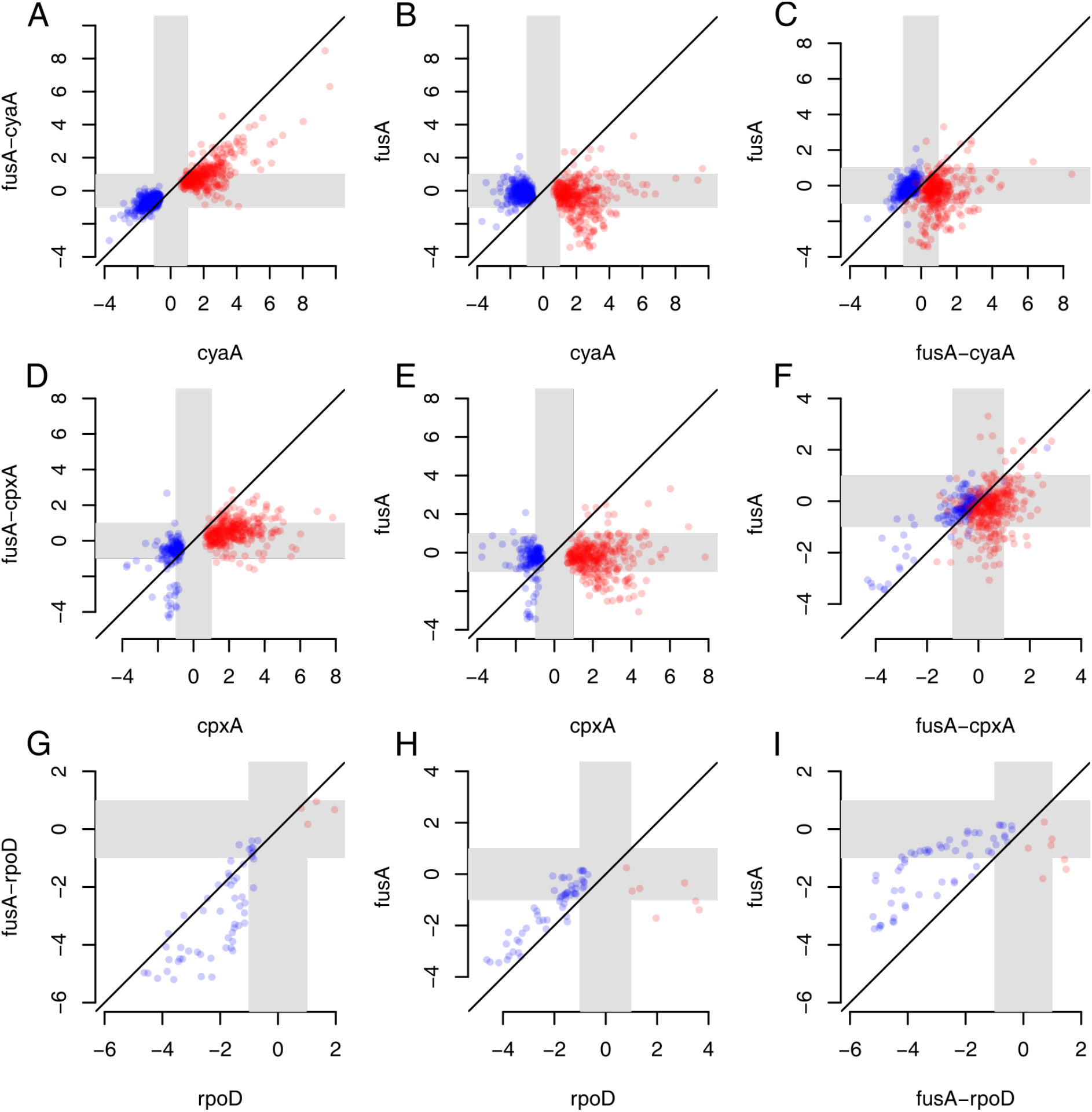
**Effect of second site mutations on gene expression and dependence on EF-G.** Scatter plots comparing logi fold changes of differentially expressed genes among mutants. Mutants are referred to by their gene names for brevity. Grey zones indicate the region between the log2 fold changes of +1 and −1 (corresponding to fold changes of 2 and 0.5), and thus highlights the region of low / no fold change. Red points show genes up-regulated and blue points show genes down-regulated in the relevant second site single mutant. (A-C) Fold changes of genes differentially expressed in the CyaA^N600Y^ mutant were compared with the FusA^A608E^-CyaA^N600Y^ and FusA^A608E^ mutants. (D-F) Fold changes of genes differentially expressed in the CpxA^F218Y^ mutant were compared with the FusA^A608E^-CpxA^F218Y^ and FusA^A608E^ mutants. (G-I) Fold changes of genes differentially expressed in the RpoD^L261Q^ mutant were compared with the FusA^A608E^-RpoD^L261Q^ and FusA^A608E^ mutants.

We saw that this was the case with the CpxA^F218Y^ and FusA^A608E^-CpxA^F218Y^ mutants as well (Fig. 6D-F). As a result of this, the number of differentially expressed genes in the FusA^A608E^-CyaA^N600Y^ or FusA^A608E^-CpxA^F218Y^ mutants is lesser than that in the CyaA^N600Y^ or CpxA^F218Y^ mutants (Fig. 5A). The effect of the mutation in *fusA* on the fold changes of genes seems to be more extreme in the case of the mutation in *cpxA* than *cyaA*, since FusA^A608E^-CpxA^F218Y^ does not correlate as well with CpxA^F218Y^ as does FusA^A608E^-CyaA^N600Y^ with CyaA^N600Y^ (Fig. 5B). These observations bear in part from the fact that the fold changes of genes differentially expressed in the CyaA^N600Y^ and CpxA^F218Y^ mutants are opposite to the fold changes of these genes in the FusA^A608E^ mutant (Fig. S23A and B), even though they were not classified as differentially expressed in FusA^A608E^. In other words, a gene that is up-regulated in CyaA^N600Y^ / CpxA^F218Y^ displays a mild negative fold change in FusA^A608E^; whereas one that is down-regulated in the former shows a slight positive fold change in the latter (Fig. S23A and B).

We notice that the effect of FusA^A608E^ on RpoD^L261Q^ is opposite to that on CyaA^N600Y^ / CpxA^F218Y^. Genes down-regulated in the RpoD^L261Q^ mutant were further down-regulated in the FusA^A608E^-RpoD^L261Q^ mutant (Fig. 6G-I). In this case, many down-regulated genes in RpoD^L261Q^ or FusA^A608E^-RpoD^L261Q^ were also down-regulated in the FusA^A608E^ mutant (Fig. 6 H and I). This bears from the fact that genes down-regulated in the RpoD^L261Q^ mutant are also down-regulated in the FusA^A608E^ mutant, thus leading to more extreme down-regulation in the FusA^A608E^-RpoD^L261Q^ mutant (Fig. S23C).

Unfortunately we do not have information of this sort for the FusA^A608E^-TopA^S180L^ mutant since we did not have the corresponding second site single mutant. However, it is possible that the interaction between the mutation in fusA and the mutation in *topA* must be extreme since we were unable to construct the *topA* single mutant.

Thus the mutation in a translation elongation factor has a large effect on the transcriptional state of the cell, beyond that indicated by threshold-dependent calls of differential expression, presumably through feedback from levels of partially folded proteins. The second-site mutations display epistasis with the FusA^A608E^ mutation, as manifested in their transcriptomes.

### Genes with known roles to play in aminoglycoside resistance are mis-regulated in the mutants

Are the mechanisms of kanamycin resistance different or common across our set of mutants? To understand this, we looked at the kinds of genes differentially expressed in these mutants using gene ontology (GO) or known transcription factor-target interactions to guide us.

Common trends in terms of shared gene functions are outlined in (Fig. 5C) and a more detailed list is provided in Fig. S20. In general, we found several metabolism related genes mis-regulated in the mutants. Mis-regulated genes with known roles in aminoglycoside resistance include genes involved in oxidative phosphorylation, protein folding and motility.

Genes involved in oxidative phosphorylation are down-regulated in the FusA^A608E^ mutant. Oxidative phosphorylation produces reactive oxygen species (ROS), as a by-product, which is thought to be involved in antibiotic mediated killing (Kohanski et al., 2007). A functional proton motive force generated by oxidative phosphorylation is required for aminoglycoside uptake (Taber et al., 1987). Furthermore, the components of the electron transport chain (ETC) tend to be Fe-S proteins and are membrane associated. Mistranslation of membrane associated proteins induced by aminoglycosides, and hence their misfolding could affect the integrity of the cell membrane and result in hydroxyl radical mediated cell death (Kohanski et al., 2008). Misfolded versions of these proteins could also release Fenton reactive Fe^2+^, which in turn could again result in hydroxyl radical generation (Kohanski et al., 2007). Thus, there are many ways in which down-regulating genes involved in oxidative phosphorylation, as in the FusA^A608E^ mutant, can alleviate the lethal effects of kanamycin.

Apart from these genes, genes associated with enterobactin biosynthesis or iron homeostasis are known to affect intracellular ROS levels (Méhi et al., 2014). It is possible that the up-regulation of these genes in the FusA^A608E^, FusA^A608E^-CpxA^F218Y^ and FusA^A608E^-TopA^S180L^ mutants could reduce the production of ROS in the cells via sequestration of free Fe^2+^ and hence contribute to resistance.

The downregulation of oxidative phosphorylation and the up-regulation of siderophore metabolism genes are in line with the hypothesis of oxidative damage mediated cell death in the presence of antibiotics. While this theory is fiercely disputed, it is possible that ROS aggravate the effect of antibiotics, if not dominate it. For example, Ling et al., show that aminoglycoside induced protein aggregation is prevented by hydrogen peroxide quenchers (Ling et al., 2012). Dealing with ROS might result just in that extra protection that cells need in the presence of aminoglycosides.

ROS is a double edged sword. While it can damage cellular macromolecules and confer stress, the outcome of this stress could also result in an increase in mutagenesis (Kohanski, DePristo & Collins, 2010). Thus if these mutants do indeed reduce oxidative stress, it is quite possible that they will slow down further adaptation to kanamycin by reducing the occurrence of resistance conferring mutations. This however remains to be tested.

We see a strong and consistent down-regulation of motility associated genes, in all except the *cyaA* mutants. Using a transposon mutagenesis screen Shan et al., show that the loss of these genes results in decreased persister formation in aminoglycosides (Shan et al., 2015), and thus their down-regulation in the mutants is contrary to our expectation. However these genes are up-regulated in the CyaA^N600Y^ and FusA^A608E^-CyaA^N600Y^ mutants and there they could contribute to resistance. Notably, motility genes are not found in the list of knockouts sensitive to aminoglycosides provided by Tamae et al. (Tamae et al., 2008), or in the list of loci that significantly affect susceptibility to aminoglycosides in the transposon insertion screen performed by Girgis et al (Girgis, Hottes & Tavazoie, 2009).

Resistance to aminoglycosides can be provided by genes involved in protein transport/ folding/ secretion as these could help refold misfolded proteins (Goltermann, Good & Bentin, 2013). However, we see a down-regulation of these genes in the FusA^A608E^-RpoD^L261Q^, FusA^A608E^-CpxA^F218Y^ and FusA^A608E^-TopA^S180L^ mutants, and this is contrary to our expectation.

## Conclusions

We saw that mutations that modify global transcriptional regulatory networks increase the resistance of the kanamycin conferring mutation in the translational elongation factor EF-G. These “second-site” mutations resulted in large changes in gene expression and displayed epistatic interactions with the mutation in EF-G, which itself drove expression changes of many genes. We show that these second site mutations reduce the activities of CyaA (adenylate cyclase) and TopA (Topoisomerase I), and increase the activity of CpxA. Further evolution of an EF-G mutant in higher concentration of kanamycin suggested CpxA as the next target for an increase in resistance, with many high frequency mutations located in the helix-I domain of this protein. Although the activity of the mutated adenylate cyclase is reduced in the CyaA mutant that we isolated, many CRP targets are unexpectedly up-regulated and this contradictory behaviour needs further investigation. We suggest that FusA^A608E^-TopA^S180L^ results in a reduction in the function of Topoisomerase I. Many genes with known roles in aminoglycoside resistance, for example genes involved in oxidative phosphorylation and enterobactin metabolism, were mis-regulated in these mutants thus pointing to possible mechanisms of resistance.

## Accession Numbers

RNA-seq data can be found on the NCBI Gene Expression Omnibus database (Edgar, Domrachev & Lash, 2002; Barrett et al., 2013) with the GEO Series accession number GSE82343 (http://www.ncbi.nlm.nih.gov/geo/query/acc.cgi?acc=GSE82343). Deep-sequencing data of populations in the 15-kan evolution experiment can be found with the accession number SRP076371 and deep-sequencing data of some strains can be found with the accession number SRP087477 from the NCBI Sequence Read Archive.

## Acknowledgements

We thank Parul Singh, NCBS, for providing *E. coli* MG1655 Δ*cyaA* and Δ*crp* strains and for sharing the transcriptome data (unpublished) of these strains. We thank Sunil Laxman, NCBS, for his guidance with cAMP estimation experiments. We thank Reshma T. V. and Revathy Krishnamurthy, NCBS, for help with some experiments. We thank Charles Dorman and Aoife Colgan, Trinity Collge Dublin, for their help with the chloroquine gel assay. Next-generation sequencing services were provided by C-CAMP and Genotypic, India.

